# High laboratory mouse pre-weaning mortality associated with litter overlap, advanced mother age, small and large litters

**DOI:** 10.1101/2020.02.25.953067

**Authors:** Gabriela Munhoz Morello, Jan Hultgren, Sara Capas-Peneda, Marc Whiltshire, Aurelie Thomas, Hannah Wardle-Jones, Sophie Brajon, Colin Gilbert, I. Anna S. Olsson

## Abstract

High and variable pre-weaning mortality is a persistent problem among the main mouse strains used in biomedical research. If a modest 15% mortality rate is assumed across all mouse strains used in the EU, approximately 1 million more pups must be produced yearly to compensate for those which die. A few environmental and social factors have been identified as affecting pup mortality, but optimizing these factors does not cease the problem. This study is the first large study to mine data records from 219,975 pups from two breeding facilities to determine the major risk factors associated with mouse pre-weaning mortality. It was hypothesized that litter overlap (i.e. the presence of older siblings in the cage when new pups are born), a recurrent social configuration in trio-housed mice, is associated with increased newborn mortality, along with high mother age, large litter size, as well as a high number and age of older siblings in the cage. The estimated probability of pup death was two to seven percentage points higher in cages with compared to those without litter overlap. Litter overlap was associated with an increase in percentage of litter losses of 19% and 103%, respectively, in the two breeding facilities. Increased number and age of older siblings, high mother age, small litter size (less than four pups born) and large litter size (over 11 pups born) were associated with increased probability of pup death. Results suggest that common social cage configurations at breeding facilities are dangerous for the survivability of young mouse pups. The underlying mechanisms and strategies to avoid these situations should be further investigated.

## Introduction

High pre-weaning mortality of laboratory mice is a major welfare and economic problem affecting mouse breeding at academic and industrial laboratories worldwide. Previous studies report pup mortalities from less than 10% [1] to as high as 49% [2] for C57BL/6 mice, one of the most commonly used mouse strains. Despite the general ongoing effort to reduce the number of animals in research and improve their welfare according to the 3R principle for research[3], high pre-weaning mortality rates persist and very little systematic research has been done to identify causes of poor survival. Data from experimental and observational studies conducted by the authors of this work at different breeding facilities in three different countries revealed that 32% of 344 litters (retrospective analysis, Germany[4]), 33% of 55 litters (experimental data, U.K.[5]), and 18% of 510 litters (experimental data, Portugal[6]) were completely lost, with the overall mortality varying from 25[6] to 52% in trio-bred mice[5] in the experimental studies. If a modest level of 15% mortality is assumed across all mouse strains, at least 1 million more mice must be produced every year just in the European Union (EU) to compensate for pups that die before they can be used in science (estimate based on the number of mice used yearly in research in the EU; European Commission 2020 [7]). Such losses are contrary to the 3R principle that is now explicit in EU legislation[8] and incur extra breeding costs of €5-8 million yearly. Several environmental, management and behavioural factors have been linked to pup mortality, such as thermal environment of the cage, level of parental care, mother age, litter size, provision of nest material, and cage manipulation [5,9–13], but manipulating these factors has not as yet eliminated the mortality problem. Recently, we identified the presence of older litter mates in the cage when a new litter is born (litter overlap) as a major factor affecting pup survival [5]. In a study with 55 litters of C57BL/6 mice (n=521 pups) housed in trios [5], a 2.3 fold increase was found in litter loss in cages where older littermates were present, compared to trio cages with no older littermates. Litter overlap happens in both trio (two adult females and one male) and pair (one adult female and one male) housing, which are the most common configurations in mouse breeding. Although litter overlap is more frequent in trios due to the presence of two breeding females, the number of pups weaned per litter is not reduced in trios compared to pairs, while trios wean more pups per cage [14]. One possible reason for this is that litter overlap in pair cages affects pup mortality more severely as compared to trio cages. In pair cages, litter overlap occurs when the only female of the cage gives birth before weaning her previous litter. In these cases, the age gap between litters becomes large, which might be especially detrimental to pup survival.

Previous research into factors affecting laboratory mouse reproduction used primarily experimental study approaches, where the sample size was small and animal management and data collection differed from standard practice in a breeding facility. With the increasing use of breeding management software, it is now possible to use much larger datasets representing the reality of practical laboratory mouse breeding. In this study, a dataset of 219,975 pups was analysed from two different collaborating breeding facilities in the UK (58,692 and 161,283 pups), by modelling the risk of a newborn mouse dying as a function of the age and number of older littermates, as well as of mother age. It was hypothesized that litter overlap is a recurrent social configuration and that the risk of pup mortality increases with litter overlap, high mother age, large litter size, as well as a high number and age of older siblings in the cage.

## Material and methods

### Data retrieval

Historical mouse breeding data was provided by two collaborating facilities. Therefore, this study did not involve any type of animal manipulation, observation, or use. Mouse breeding in the collaborating facilities was performed in line with the UK Animals (Scientific Procedures) Act of 1986.

Mouse breeding data were made available by two collaborating breeding facilities (C1, the Babraham Institute and C2, the Wellcome Sanger Institute). Historical production data were downloaded directly from their breeding management software (MCMS, Mouse Colony Management System, the Wellcome Sanger Institute Data Centre). C1 provided data from January 2014 to October 2018, and C2 provided data from January 2010 to March 2019. The datasets contained information on litter identity, breeding adults’ identities, date of birth, date of death, number of pups born and number of pups weaned.

### Animals, housing, and management

The original dataset contained a total of 34,949 C57BL/6 litters and 219,975 pups. All mice were housed in trios (two females and one male) in individually ventilated cages (IVC). Details on animals, housing, and management are shown in Table 1.

**Table 1.** Animal, housing, and management characteristics for the collaborating animal facilities.

|  | **The Babraham Institute, C1** | **The Wellcome Sanger Institute, C2** |
| --- | --- | --- |
| **Mouse Strain** | C57BL/6BabR | C57BL/6NTac |
| **Housing configuration** | Trios | Trios |
| **Type of cages** | IVC<sup>a</sup> Tecniplast GM500, transparent polysulphone | IVC<sup>a</sup> Tecniplast GM500, transparent polysulphone and Tecniplast Sealsafe 1284L |
| **Ventilation rate** | 65 to 75 air changes/hour | 60 air changes/hour |
| **Air handling unit** | Tecniplast DGM80 and DGM160 | Tecniplast TouchSLIMLine™ |
| **Bedding** | 5 mm deep soft-wood-flake bedding (ECO6, Datesand group, Manchester, UK) | 175 g of Aspen Chips (B&K Universal Ltd, Peninsula Plaza, Singapore) |
| **Nest** | 7 g of white paper rolls (Enrich-n'Nest, Datesand group, Manchester, ENG, UK) | 25 mm square Nestlet (Datesand group, Manchester, UK) derived from pulped cotton virgin fibre |
| **Enrichment** | Tecniplast Mouse Pouch Loft, a second level flooring within the cage | One cardboard bio-tunnel per cage |
| **Water** | *Ad libitum*, sterilized through reverse osmosis and provided through automatic drinking valves (Edstrom A160/QD2, Avidity Science LLC, Waterford, WI, USA) | *Ad libitum*, triple filtered, provided in flash sterilized acrylic bottles with stainless steel drinking caps |
| **Food** | *Ad libitum* in the form of standard 9.5 mm diameter dry pellets (CRM(P), Special Diets Services, Witham, Essex, UK) | *Ad libitum* in the form of standard 10.5 mm diameter dry pellets (SAFE R03-10, Augy, France) |
| **Cage change routine** | Cages changed once every second week; cages with large litters sometimes cleaned every week | Cages assessed once a week and changed if needed |
| **Room temperature (target)** | 20°C to 21°C | 19°C to 23°C |
| **Room relative humidity (target)** | 50% | 45% to 65% |
| **Light schedule** | 12 hours light (7:00-19:00) and 12 hours dark | 12 hours light (7:30-19:30) and 12 hours dark |
| **Weaning age (target)** | 21 days | 19 to 23 days; male pups were euthanized before weaning |
| **Breeding start age (target)** | 8 to 9 weeks | 6 to 9 weeks |
| **Retirement age (target)** | 24-32 weeks; longer if productive | 24 weeks, or after 3 poor litters, or after 5 or 6 successful litters |
| **Pup counting routine** | Pups are counted once between their birth and day 7 pp with minimal handling | Pups are counted whenever cages are cleaned. Pups less than five 5 d old are left undisturbed (not counted) |

<sup>a</sup>IVC=Individually ventilated cage

### Statistical analysis

Data were collected from 58,692 pups in 9,261 litters and 161,283 pups in 25,688 litters from C1 and C2, respectively. Required information was retrieved using Scilab (version 6.0.1, Scilab Enterprises, Rungis, France), resulting in one data line per pup. A total of 11% (C1) and 21% (C2) of the data provided was excluded, mainly due to incongruent data records, implausibly large litters (more than 13 pups, unless confirmed as correct), unreliable information on number and age of older pups in the cage, or missing information. Male pups at C2 euthanized at day 7 pp or later were coded as surviving. Litters with males euthanized before day 7 pp were excluded.

Pup mortality before weaning was coded as 0 (survived) or 1 (died) and used as the dependent variable. Environmental and social factors were considered as risk factors for pup death. Independent variables representing environmental factors considered for analysis were Collaborator (C1 or C2), Season (Winter, Spring, Summer, Fall), Month (as an alternative predictor to Season), Weekday, and Year, while independent variables for social risk factors included Mother Age (continuous), Father Age (continuous), Litter Size (number of pups born; continuous), litter Overlap (whether or not older siblings were present at the time of birth of the focus litter; no or yes), Sibling Number (number of older pups in the cage at the time of birth of the focus litter; continuous) and Sibling Age (age of the older siblings; continuous).

The risk of pup death was modelled by mixed logistic regression, using the GLIMMIX procedure in SAS (2018 University Edition, SAS Institute Inc., Cary, NC, USA). Multicollinearity among independent variables was checked by using the variance inflation factor (VIF) and regressing each independent variable on the others. As a consequence, Year, Month, and Father Age were excluded from the analysis.

Data of C1 and C2 were combined together and two separate models were constructed; one model using data for all pups, including those with and without litter overlap, and another model for pups with litter overlap only. In both models, litter identity was included as a random effect to account for clustering. The models were built by adding one independent variable of interest (Mother Age, Litter Size, Overlap (first model) or Sibling Number and Sibling Age (second model)) at a time in a stepwise process with bidirectional elimination. Independent variables with P ≤ 0.05 were kept in the model. Weekday, Season, and Collaborator were then tested one at a time as confounders, followed by possible interactions and higher order terms. Least-squares means of Weekday, Season, and Overlap were examined and compared among different variable levels.

## Results

The percentage of pups dying before weaning was 39% at C1 and 14% at C2, while the mean number of Litter Size was 7.6 pups born/litter in both collaborators. In 42% of the C1 litters and 78% of the C2 litters no pups died. The percentage of litters with at least 90% death rate was 28% at C1 and 9% at C2. Approximately 50% and 57% of the litters were born with the presence of older siblings in the cage (litter overlap) in C1 and C2, respectively.

The first model (all pups) contained the variables Collaborator, Season, Weekday, Mother Age, Litter Size and Overlap. The second model (pups with litter overlap) contained variables Collaborator, Season, Weekday, Mother Age, Litter Size, Sibling Number, and Sibling Age. Collaborator interacted significantly with all the independent variables in both models, except Mother Age in the second model. Therefore, results for C1 and C2 are presented separately. Sibling Number interacted significantly with Sibling Age, while Litter Size affected pup death probability in a quadratic fashion in both models. Model details are available in S1 and S2 Tables.

The estimated probability of pup death was seven (C1) and two (C2) percentage points higher (P < 0.01) in cages with the presence of older siblings compared to cages without an older litter (Fig 1A and 1B). At C1, 31% of the overlapped and 26% of the non-overlapped litters had a total litter loss (all pups dying, Fig 1C), whereas at C2, the corresponding figures were 12% and 6% (Fig 1D).

**Fig 1.**
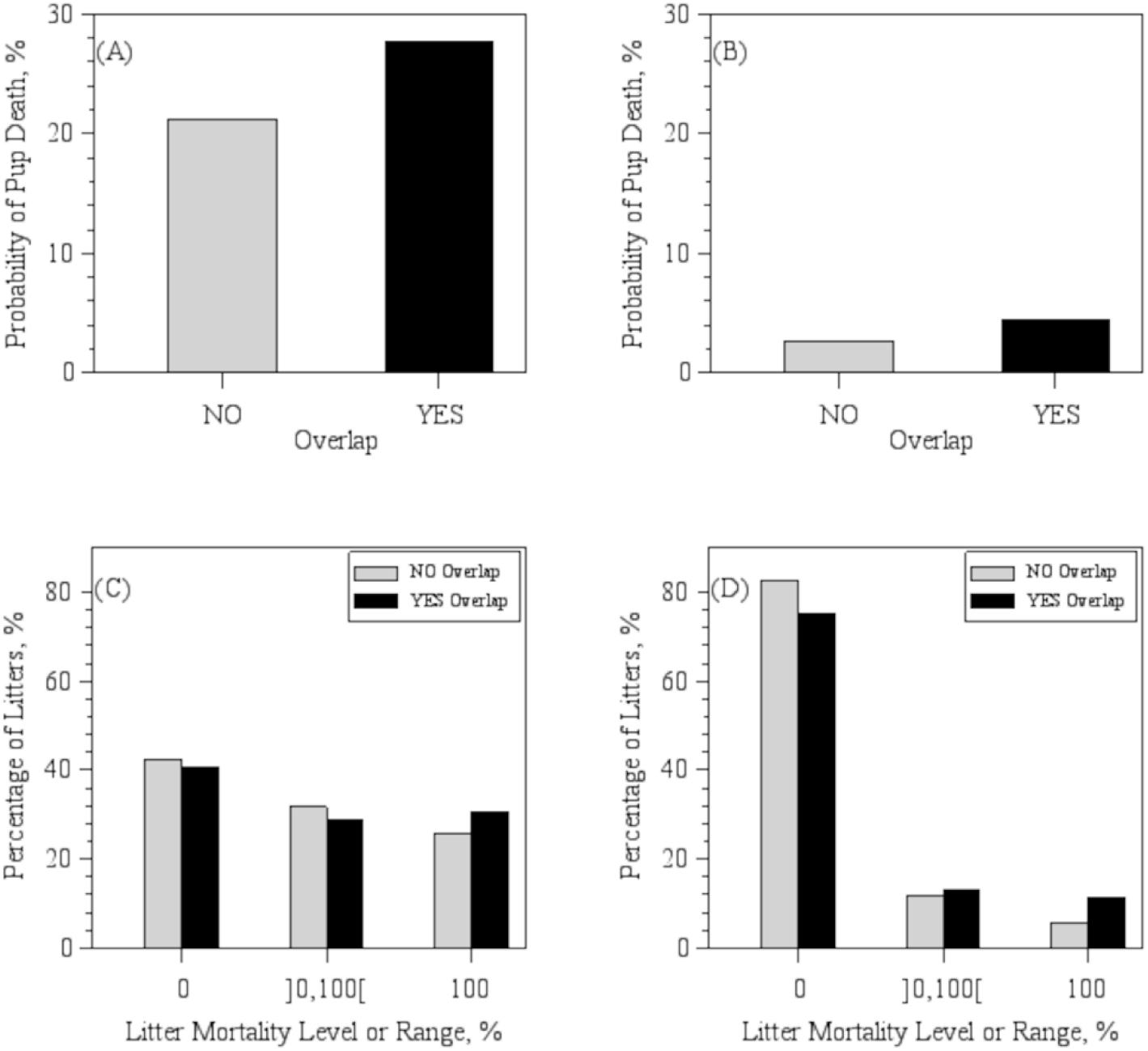
Probability of pup death and litter mortality distribution. Probability of pup death in litters without (NO) or with (YES) the presence of older siblings in the cage (litter overlap) at (A) C1, the Babraham Institute, and (B) C2, the Wellcome Institute, based on least-square means. Percentage of litters born with litter overlap by category of pre-weaning mortality at (C) C1 and (D) C2, based on raw data. Numbers within brackets in the x-axis designate lower (left side) and upper (right side) limits of mortality range. An open bracket next to a number designates a non-inclusive limit.

The predicted probability for a pup to die as a function of Mother Age, Litter Size, Sibling Age, and Sibling Number is illustrated by Figs 2 and 3. Increased Mother Age, Sibling Number, and Sibling Age were associated (P < 0.01) with an increase in the probability of pups dying at both C1 and C2 (Figs 2 and 3).

**Fig 2.**
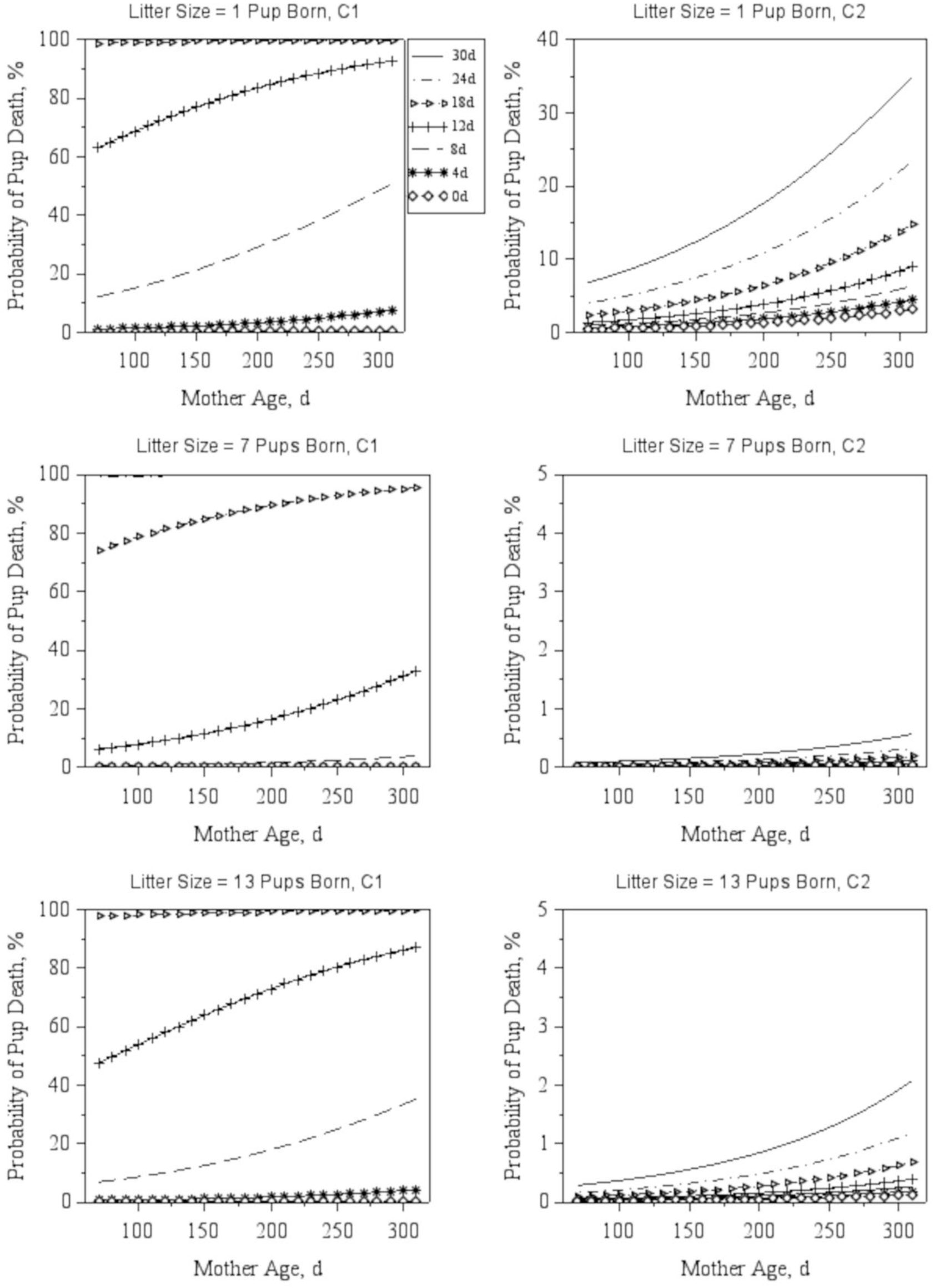
Probability of pup death by Mother Age and Sibling Age. Predicted probabilities (least-squares means) of a pup to die as a function of Mother age for three distinct levels of Litter Size (number of pups born), at C1, the Babraham Institute and C2, the Wellcome Sanger Institute. Each line corresponds to predictions for a specific value of Sibling Age, as depicted in the legend next to the top left graph. Predictions were obtained while assuming six older pups in the cage, the most recurrent Weekday (Thursday) and the most common Season (Spring) in the combined dataset.

**Fig 3.**
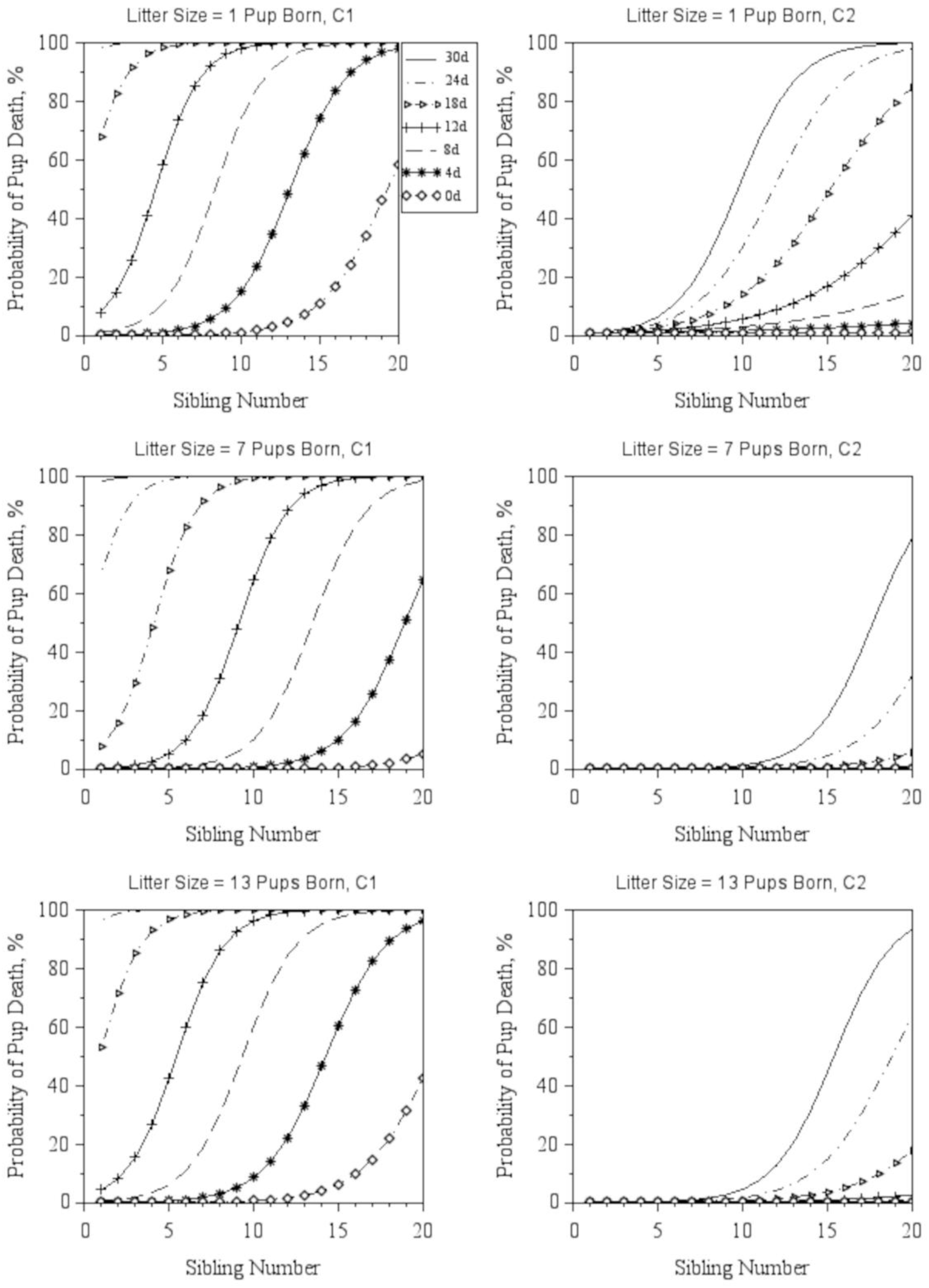
Probability of pup death by Sibling Number and Sibling Age. Predicted probabilities (least-squares means) of a pup to die as a function of Sibling Number (number of older pups in the cage at the birth of the focal litter) for three distinct levels of Litter Size (number of pups born), at C1, the Babraham Institute and C2, the Wellcome Sanger Institute. Each line corresponds to predictions for a specific value of Sibling Age, as depicted in the legend next to the top left graph. Predictions were obtained while assuming six older pups in the cage, the most recurrent Weekday (Thursday) and the most common Season (Spring) in the combined dataset.

Pup death was associated with Litter Size in a quadratic fashion in both collaborators (P < 0.01). For all combinations of Mother Age and Sibling Age, the risk of death was nearly at its minimum around a litter sizes of 6-10 pups (mean litter size at birth for overlapped data: 7.3 ± 2.6 pups). Although Weekday and Season were not added to the models as variables of interest, these factors turned out to be confounders to the models (P < 0.01). The probability of death consistently decreased towards the end of the week at both collaborators, while the effect of Season lacked a consistent pattern between collaborators (available in S1 Fig). C1 was able to provide records on cage cleaning dates per Weekday for the period of 17 months (April 2018 to November 2019) for the colony. Cage change events also peaked in the beginning of the week and were lower towards the end of the week (available in S2 Fig).

## Discussion

Neonatal mortality is a large problem in laboratory mouse breeding and improving pup survival is a key to improve the efficiency and sustainability of producing laboratory mice while complying with the 3R principle[8]. Here, we present the first large scale study of management, environmental and animal factors affecting pup survival based on data from over 200,000 C57BL/6 pups born during a period of 5 to 10 years in two large mouse breeding facilities. The study indicates that litter overlap, a social configuration which frequently occurs in trio-breeding cages, results in a 30 to 60% increase in the probability of neonatal pup death. The higher the number and age of older siblings in the cage, the greater is the risk of neonatal pup death. The probability of pups to die in the presence of older siblings was also affected by mother age and litter size.

Irrespective of any mortality risks identified, the average litter mortality rates obtained both at C1 (39%) and C2 (14%) were higher than what was previously reported for C57BL/6J mice (8% pre-weaning mortality[15] with pup counting at weaning and 3% mortality at three days pp[16] with pup number obtained from video-records). Litter mortality was higher in C1 compared to C2 and no differences were found between collaborators in the number of pups born per litter. Husbandry differences between C1 and C2 may contribute to the mortality difference between both institute as cage temperature, as well as nest and bedding amount and quality affect rodents’ breeding performance[10–13]. An additional factor may be the practice of euthanizing male pups in C2. Euthanized males after seven days of age were considered as survivals, but this is an assumption and may have led to an underestimation of mortality. Finally, any differences in accuracy of data entry may have affected the results. C1 had a more consistent and early counting of pups than C2. Thus it is possible that C1 has a better accuracy in detecting the number of pups born compared to C2 where pups born could be underestimated, considering that most of deaths happen within 48h pp and dead pups often get cannibalized by the dam, thus not seen by the caretakers. Productivity differences either in pup survivability or in the actual number of pups born due to the distinct mouse sub-strains between C1 and C2 could also underlie the differences in overall mortality found between C1 and C2.

The found higher mouse pre-weaning mortality in trios with overlapped litters, i.e. litters born when an older litter was present, compared to non-overlapped litters is in agreement with previous experimental findings. In outbred mice derived from the C57BL/6J, BALB/c, and DBA/1J strains, Schmidt et al. (2015)[17] found pup mortality to increase with increasing age gap between the litters sharing a cage at a specific time. Understanding why being born into a cage with an older litter is so dangerous requires information about events around pup death and the condition of dead and dying pups. Past research have frequently associated pup mortality with infanticide[17,18], assuming that cannibalized pups were killed before they were eaten. However, previous behavior studies conducted by the authors of this study revealed that infanticide precedes less than 15% of the cannibalism events[5,19] and that pups die primarily from other causes than direct killing. Litter asynchrony, which often leads to overlap, is likely to increase unequal competition for access to milk and parental care, trauma caused by trampling and stepping of newborns by the adults or the older siblings, and problems related with increased cage stocking density.

Early access to milk is essential for the survival of newborn pups. Measurements of pup energy losses due to metabolism between nursing bouts, extrapolated for a period of 24 hours, revealed that if pups did not receive milk during their first day of life, they would lose approximately 8% of their birth weight[20], which would likely reduce their chances of survival. The presence of older and consequently heavier, more developed and more mobile pups in the cage may have interfered with the access of the newborns to milk in general and specifically to steal the iron-rich milk that is present in higher concentrations during the first week of lactation[20]. Also, older and heavier siblings may be able to displace light newborn pups more easily from the dam’s nipples, as compared to younger siblings. This could partly explain the interaction found in this work between the Number and the Age of Older Siblings in the cage, affecting pup death probability. The presence of two litters in the cage has been demonstrated to increase parturition duration and affect parental behavior. Adults in trio cages with two litters were observed to care for their newborn pups a total of 20% less time (all the three adults together) than adults with one single litter in the cage[5], while parental investment is known to improve the chances of survival of young mice. In fact, C57BL/6 females which lost their litters entirely have been found to spend more time outside the nest and invested less time in building the nest prior to parturition[21], while the presence of males in cages with breeding females (CD-1) has been demonstrated to increase pup survival by facilitating maternal behavior[22]. Thus, reduced parental care in cages with more than one litter can be one of the mechanisms through which pup survivability is reduced in the presence of an older litter.

Most often, when there are two females sharing a cage, they also share the same nest, and younger lighter pups get clustered together with the older, heavier, and more mobile pups. Data on post-mortem inspection performed in 324 C57BL/6J pups found dead, by the authors of this study, revealed that 24% of the pups had some kind of traumatic lesion, including bite wounds and bruises[23].

Higher stocking density leads to increased humidity and gas concentration in the air, with ammonia reaching above 150 ppm in mouse breeding cages prior to weaning[24,25]. Whereas the impact of these ammonia levels on newborn mice has not been studied, ammonia levels from 25 to 250 ppm have been demonstrated to destroy the surface layers of the trachea epithelium lining and increase the severity of rhinitis, otitis, tracheitis, and pneumonia in rats and mice[25–27]. The gas concentration problem may be aggravated by the fact that animal care-takers generally tend to avoid cleaning cages when litters were just born (to avoid pup disturbance).

### Mother age

Pup death probability increased as Mother Age increased in both collaborators. In this study, Mother Age and parity were confounded. Thus, it was not possible to distinguish effects on pup death probability of the dam’s age and its birthing experience. Decreased productivity and increased mortality in first-parity litters have been reported for a few different species[28,29], but for mice this subject remains controversial. While first-parity BALB/c and 129/Sv dams were reported to wean fewer pups per litter compared to later parity ones[30], we previously found an increase in pup survival with lower parities[5] in an experimental study conducted in C1, whereas another study did not find any significant differences in pup loss between first- and later parity C57BL/6 or BALB/c dams[4].

The results for Mother Age are in agreement with those from Tarín et al. (2005)[9], who found increased pup mortality and incidence of litters with at least one cannibalized pup with increased parity. Tarín et al. (2005)[9] also compared breeding performance between mothers (F1 of C57BL/6JIco × CBA/JIco) who started their reproductive life at age 70 d (young) and 357 d (old). The authors found no differences between mother age group on pre-weaning mortality and litter size both at birth and at weaning, but reported that young mothers produced F2 litters with higher expectation of survival and body weight than those of old mothers.

### Number of pups born

Pup death probability was higher in either small or large litters (less than four or more than 11 pups born). The reduced survivability in small litters is in agreement with previous reports for C57BL/6[5] and F1 hybrid (C57BL/6JIco × CBA/JIco) mice[9]. This may be related to the amount of parental care. Ehret and Bernecker (1986)[31] demonstrated that early pup vocalization, which gradually increases in frequency after birth, is essential to maintain maternal attention at high levels, which leads to improved pup weight gain, as compared to pups from mothers which were unable to hear them. Therefore, it is possible that a small newly born litter does not emit sufficient vocal cues to ensure sufficient maternal care. Rat litters[32] with one single pup were found to perform only about 10% and 5% of the suckling stimuli performed by litters of 10 and 22 pups. As a consequence, the milk yield of dams (estimated based on adjusted measures of the pups’ daily weight gain) raising single pups was only −0.4% to 7.0% of those raising 10 pups, which led one-pup litters to have the lowest growth rate. More than half of the one-pup litters did not show any weight gain in the first five days pp. From an evolutionary perspective, a small litter is less worth investing in than a larger litter: Maestripieri and Alleva (1991) [33] demonstrated that CD-1 dams of large litters (eight pups) spent more than twice as much time displaying litter defense behaviors against intruder males than dams of small litters (four pups). The increase in pup death probability found in litters of 12 pups and above, on the other hand, may be a result of increased sibling competition for access to milk, as discussed above, and also may represent a ceiling in milk production capacity by the mothers[20,34,35].

### Weekday and season

In both collaborators, there seemed to be a decrease the probability of pup death towards the end of the week, possibly associated with the timing of cage changes, a management routine which affects the mice as well as the accuracy of mortality detection. In C1, which provided records on cage cleaning dates, these closely mirrored the pattern of pup death probability. To reliably count the number of pups, the cage must be opened and animals moved, something that often only happens at cage cleaning when manipulation is unavoidable. Mortality is therefore likely to be more accurately detected for litters born on cage changing days. For example, an eight-pup litter born on a Tuesday with cage cleaning schedule for the same day will be recorded as an eight-pup litter. If two of these pups die in the following 24 hours, this litter’s pre-weaning mortality will be recorded as being 25% at weaning. A similar litter born on a Saturday with two pups dying on Sunday, and subsequently cannibalized, will be recorded at the Tuesday cage change as a litter with six pups born with no pre-weaning deaths.

Still, the mouse disturbance hypothesis cannot be disregarded. If pup mortality is affected by cage change, the same pattern would be expected in cage change frequency as in pup death probability per Weekday (of birth). Reeb-Whitaker et al. (2000)[1] found a higher pup mortality in cages with weekly changes than those changed once every two weeks. Cage change requires that mice are moved from the dirty to a clean cage, an event that triggers a stress response evidenced by increases in serum corticosterone[36] and general activity[37]. It is possible, therefore, that cage change interferes with parental behavior in breeding cages, which could aggravate pup mortality around those days.

### Conclusions

The present study revealed that high pre-weaning mortality in laboratory mice (C57BL/6) is associated with advanced mother age, litter overlap, the presence of a high number and age of older siblings in the cage, and a small (less than four) or large (more than 11 pups) litter. The dynamics of parental care, sibling competition for access to milk, and issues related with the number of animals in the cage may underlie the effects found in pup mortality caused by the identified risks. Future studies should address sibling competition and parental behavior in asynchronized litters.

## Ackowledgements

Dr. Luiz Henrique Antunes Rodrigues for providing technical input for the statistical modeling process. The Babraham Institute and the Wellcome Sanger Institute, UK, for providing the historical data used in this manuscript. Dr. Sofia Lamas for providing technical and scientific input.

## Supporting information

**S1 Fig.**
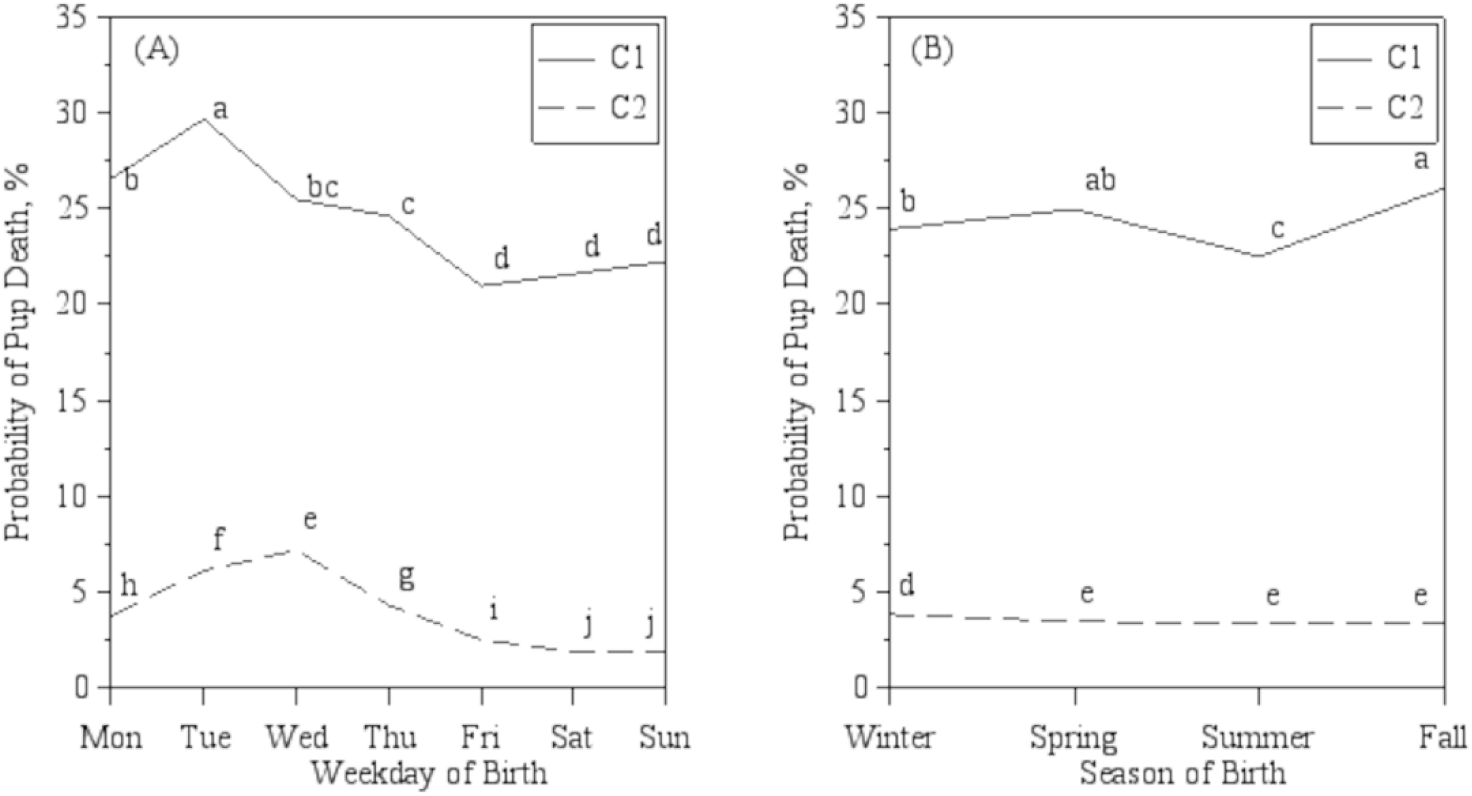
Probability of pup death by Weekday and Season. Predicted Probability (least-square means) of a pup to die as a function of (A) Weekday and (B) Season of birth at C1, the Babraham Institute, and C2, the Wellcome Sanger Institute. Data points with distinct labeled letters indicate statistical difference at 95% confidence level. Probability of a pup to die is depicted in terms of least-square means.

**S2 Fig.**
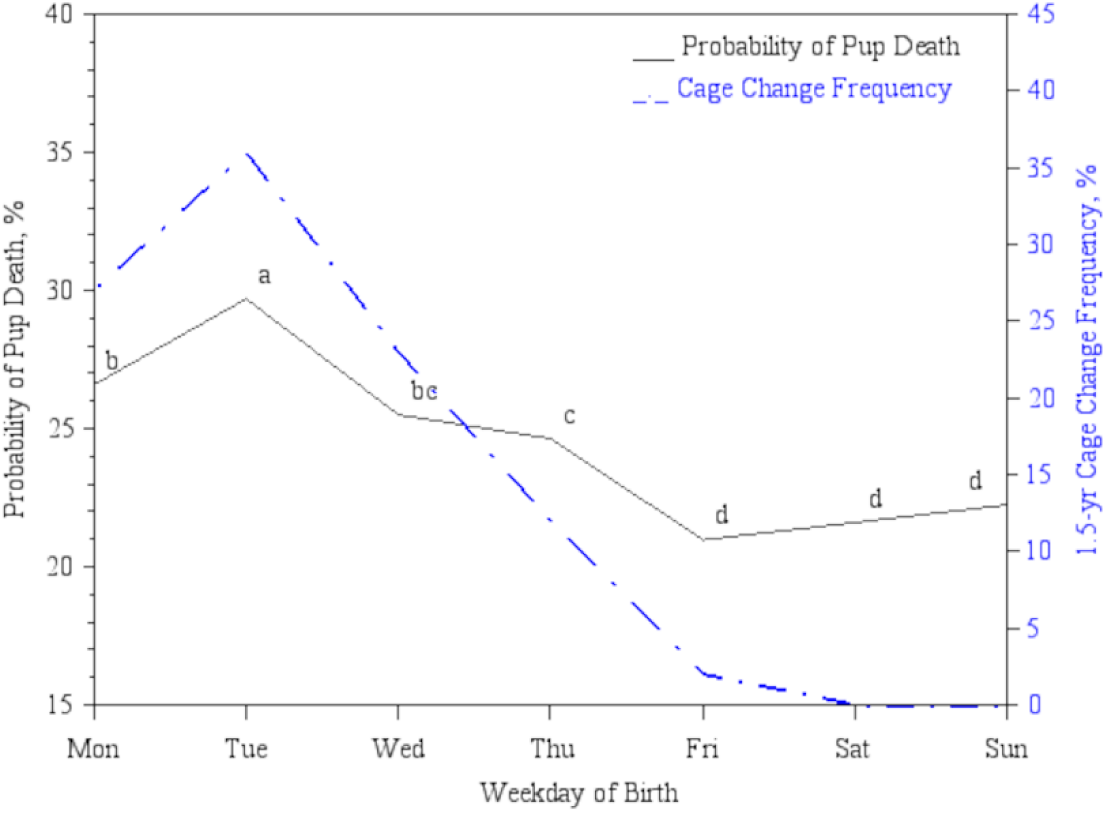
Cage change frequency and probability of pup death by Weekday. Cage change frequency and predicted probability (least-square means) of a pup to die as a function of Weekday at C1, the Babraham Institute. Data points with distinct labeled letters indicate statistical difference at 95% confidence level. Cage change frequency is depicted as the percentage per weekday of the 78 cage change episodes which happened from April 2018 to November 2019 (available data records), in the studied room of C1.

**S1 Table.** Solutions for fixed effects of the model predicting the odds of pup death fitted in the whole processed dataset.

| Effect | Estimate | Standard Error | t Value | Pr > \|t\| |
| --- | --- | --- | --- | --- |
| Intercept | -1.0546 | 0.0751 | -14.04 | <0.0001 |
| Collaborator C1 | 2.3067 | 0.1011 | 22.81 | <0.0001 |
| Collaborator C2 | 0.0000 | n.a. | n.a. | n.a. |
| Season Fall | -0.1468 | 0.0354 | -4.14 | <0.0001 |
| Season Spring | -0.1082 | 0.0360 | -3.00 | 0.0027 |
| Season Summer | -0.1234 | 0.0400 | -3.09 | 0.0020 |
| Season Winter | 0.0000 | n.a. | n.a. | n.a. |
| Weekday 1 | -0.0054 | 0.0517 | -0.11 | 0.9162 |
| Weekday 2 | 0.7230 | 0.0472 | 15.33 | <0.0001 |
| Weekday 3 | 1.2433 | 0.0463 | 26.86 | <0.0001 |
| Weekday 4 | 1.4133 | 0.0453 | 31.18 | <0.0001 |
| Weekday 5 | 0.8705 | 0.0461 | 16.88 | <0.0001 |
| Weekday 6 | 0.3000 | 0.0482 | 6.22 | <0.0001 |
| **Weekday 7** | 0.0000 | n.a. | n.a. | n.a. |
| **Mother Age** | 0.0046 | 0.0003 | 15.89 | <0.0001 |
| **Litter Size<sup>a</sup>** | -0.7104 | 0.0134 | -53.20 | <0.0001 |
| **Litter Size<sup>2a</sup>** | 0.0345 | 0.0009 | 36.73 | <0.0001 |
| **Overlap No** | -0.4969 | 0.0240 | -20.67 | <0.0001 |
| **Overlap Yes** | 0.0000 | n.a. | n.a. | n.a. |
| **Collaborator C1\* Season Fall** | 0.2607 | 0.0488 | 5.35 | <0.0001 |
| **Collaborator C1\* Season Spring** | 0.1647 | 0.0495 | 3.33 | 0.0009 |
| **Collaborator C1\* Season Summer** | 0.0435 | 0.0541 | 0.80 | 0.4210 |
| **Collaborator C1\* Season Winter** | 0.0000 | n.a. | n.a. | n.a. |
| **Collaborator C2\* Season Fall** | 0.0000 | n.a. | n.a. | n.a. |
| **Collaborator C2\* Season Spring** | 0.0000 | n.a. | n.a. | n.a. |
| **Collaborator C2\* Season Summer** | 0.0000 | n.a. | n.a. | n.a. |
| **Collaborator C2\* Season Winter** | 0.0000 | n.a. | n.a. | n.a. |
| **Collaborator C1\* Weekday Sunday** | 0.04295 | 0.0683 | 0.63 | 0.5292 |
| **Collaborator C1\* Weekday Monday** | -0.4495 | 0.0639 | -7.04 | <0.0001 |
| **Collaborator C1\* Weekday Tuesday** | -0.8161 | 0.0637 | -12.81 | <0.0001 |
| **Collaborator C1\* Weekday Wednesday** | -1.1965 | 0.0626 | -19.11 | <0.0001 |
| **Collaborator C1\* Weekday Thursday** | -0.6996 | 0.0632 | -11.06 | <0.0001 |
| **Collaborator C1\* Weekday Friday** | -0.3375 | 0.0648 | -5.21 | <0.0001 |
| **Collaborator C1\* Weekday Saturday** | 0.0000 | n.a. | n.a. | n.a. |
| **Collaborator C2\* Weekday Sunday** | 0.0000 | n.a. | n.a. | n.a. |
| **Collaborator C2\* Weekday Monday** | 0.0000 | n.a. | n.a. | n.a. |
| **Collaborator C2\* Weekday Tuesday** | 0.0000 | n.a. | n.a. | n.a. |
| **Collaborator C2\* Weekday Wednesday** | 0.0000 | n.a. | n.a. | n.a. |
| **Collaborator C2\* Weekday Thursday** | 0.0000 | n.a. | n.a. | n.a. |
| **Collaborator C2\* Weekday Friday** | 0.0000 | n.a. | n.a. | n.a. |
| **Collaborator C2\* Weekday Saturday** | 0.0000 | n.a. | n.a. | n.a. |
| **Collaborator C1\*Mother Age** | -0.0013 | 0.0003 | -3.81 | 0.0001 |
| **Collaborator C2\*Mother Age** | 0.0000 | n.a. | n.a. | n.a. |
| **Collaborator C1\*Litter Size<sup>a</sup>** | 0.0484 | 0.0066 | 7.37 | <0.0001 |
| **Collaborator C2\*Litter Size<sup>a</sup>** | 0.0000 | n.a. | n.a. | n.a. |
| **Collaborator C1\*Overlap No** | 0.1426 | 0.0332 | 4.30 | <0.0001 |
| **Collaborator C1\*Overlap Yes** | 0.0000 | n.a. | n.a. | n.a. |
| **Collaborator C2\*Overlap No** | 0.0000 | n.a. | n.a. | n.a. |
| **Collaborator C2\*Overlap Yes** | 0.0000 | n.a. | n.a. | n.a. |

Type III (Partial) Tests of Fixed Effects
| Effect | Num DF | F Value | Pr > F |
| --- | --- | --- | --- |
| **Collaborator** | 1 | 579.79 | <0.0001 |
| **Season** | 3 | 5.84 | 0.0006 |
| **Weekday** | 6 | 256.88 | <0.0001 |
| **Mother Age** | 1 | 511.62 | <0.0001 |
| **Litter Size<sup>a</sup>** | 1 | 2563.59 | <0.0001 |
| **Litter Size<sup>2a</sup>** | 1 | 1349.04 | <0.0001 |
| **Overlap** | 1 | 658.04 | <0.0001 |
| **Collaborator\*Season** | 3 | 13.55 | <0.0001 |
| **Collaborator\*Weekday** | 6 | 102.88 | <0.0001 |
| **Collaborator\*Mother Age** | 1 | 14.48 | 0.0001 |
| **Collaborator\*Litter Size<sup>a</sup>** | 1 | 54.30 | <0.0001 |
| **Collaborator\*Overlap** | 1 | 18.46 | <0.0001 |

n.a.=not applicable.
<sup>a</sup>Variable Litter Size was centered by its mean.

**S2 Table.**
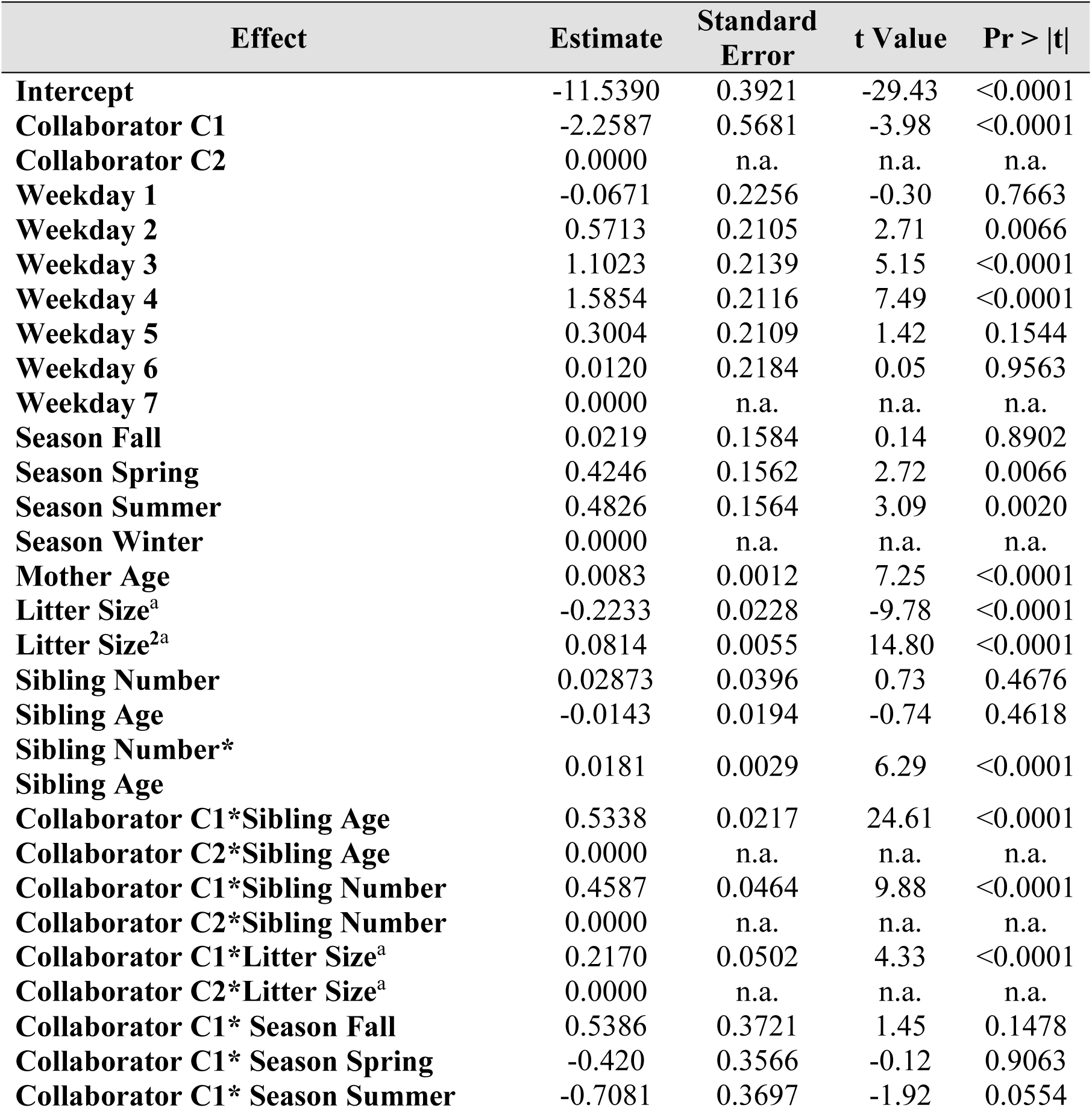
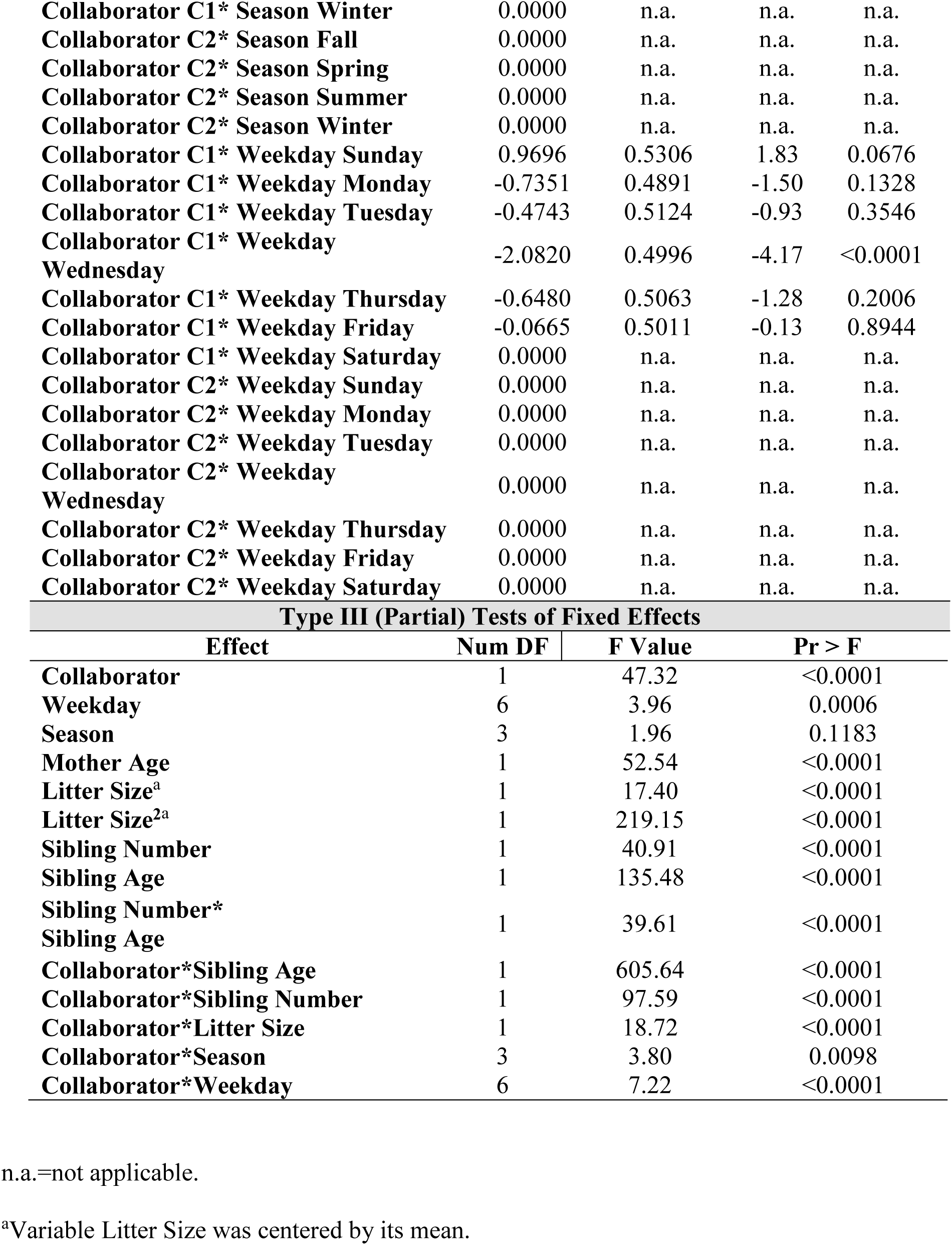
Solutions for fixed effects of the model predicting the odds of pup death fitted in the dataset containing only overlapped litters.

